# Three types of actomyosin rings within a common cytoplasm exhibit distinct modes of contractility

**DOI:** 10.1101/2024.08.08.607200

**Authors:** John B. Linehan, Alexandra Zampetaki, Michael E. Werner, Bryan Heck, Paul S. Maddox, Sebastian Fürthauer, Amy S. Maddox

## Abstract

Actomyosin rings are specializations of the non-muscle actomyosin cytoskeleton that drive cell shape changes during division, wound healing, and other events. Contractile rings throughout phylogeny and in a range of cellular contexts are built from conserved components including non-muscle myosin II, actin filaments, and crosslinking proteins. To explore whether diverse actomyosin rings generate contractile force and close via a common mechanism, we studied three instances of ring closure within the continuous cytoplasm of the *C. elegans* syncytial oogenic germline: mitotic cytokinesis of germline stem cells, apoptosis of meiotic compartments, and cellularization of oocytes. The three ring types exhibited distinct closure kinetics and component protein abundance dyanmics. We formulated a physical model to relate measured closure speed and molecular composition dynamics to ring active stress and viscosity. We conclude that these ring intrinsic factors vary among the ring types. Our model suggests that motor and non-motor crosslinkers’ abundance and distribution along filaments are important to recapitulate observed closure dynamics. Thus, our findings suggest that across ring closure contexts, fundamental contractile mechanics are conserved, and the magnitude of contractile force is tuned via regulation of ring component abundance and distribution. These results motivate testable hypotheses about cytoskeletal regulation, architecture, and remodeling.

## Introduction

The non-muscle actomyosin cytoskeleton dynamically remodels to drive numerous cell biological and developmental processes (Bear and Haugh, 2014). The abilities of an actomyosin network to rearrange, generate, and resist forces are determined by the interactions among its molecular components: flexible actin filaments (F-actin), bipolar arrays of myosin motors (non-muscle myosin II; NMMII), and non-motor F-actin crosslinkers including anillin, plastin/fimbrin, α-actinin, and filamin. Actomyosin rings are specializations of the plasma membrane-associated cortical cytoskeleton that exist in many contexts throughout animal and fungal cell physiology (Green *et al*., 2012). Constricting actomyosin rings drive cell expulsion from monolayers, healing of multi- or subcellular wounds, enucleation, and cytokinesis (Schwayer *et al*., 2016). Stable actomyosin rings can maintain cytoplasmic connections among compartments, such as in germline syncytia (see below; Haglund *et al*., 2011).

Cytokinesis, when an actomyosin ring separates the two daughter cells during division, is the context in which the functions of many cytoskeletal proteins including NMMII and anillin scaffold proteins have been defined (Green *et al*., 2012; Kamasaki *et al*., 2007; Pollard and Wu, 2010; Wu and Pollard, 2005; Schwayer *et al*., 2016; O’Shaughnessy and Thiyagarajan, 2018). Anillins are multidomain proteins that bind the plasma membrane and to structural and regulatory elements of the actomyosin cytoskeleton (Maddox *et al*., 2005; D’Avino, 2009; Piekny and Maddox, 2010; Zhang and Maddox, 2010). They contribute to the organization and therefore the effectiveness of actomyosin contractile networks (Piekny and Maddox, 2010; D’Avino, 2009; Kučera *et al*., 2021). NMMII is a motor protein that forms bipolar filamentous aggregates that crosslink and slide F-actin (Glotzer, 2005; Osório *et al*., 2019; Weeds and Lowey, 1971; Niederman and Pollard, 1975). NMMII is critical for cytokinesis in most animal and fungal cell types (Mabuchi and Okuno, 1977; De Lozanne and Spudich, 1987; Kitayama *et al*., 1997; Straight *et al*., 2003; Ma *et al*., 2012; Pollard, 2017) via its abilities to build stresses by motoring along and crosslinking F-actin (Fang *et al*., 2010; Ma *et al*., 2012; Palani *et al*., 2017; Povea⍰ Cabello *et al*., 2017; Wang *et al*., 2020; Osório *et al*., 2019). NMMII is dispensable for cytokinesis in budding yeast and slime molds likely due to the dominant mechanical contributions of cell wall deposition and cell motility, respectively (Wloka and Bi, 2012; Lord *et al*., 2005; Mendes Pinto *et al*., 2012; Bi et al., 1998; Thiyagarajan *et al*., 2015; Tolliday *et al*., 2003; Schmidt *et al*., 2002).

Consistent with the concept that various cell types share a contractility mechanism, closure speed scales with ring starting size during *C. elegans* early embryogenesis and vulval precursor cell division, as well as in the filamentous fungus *N. crassa*, such that larger cytokinetic rings close faster than smaller cytokinetic rings and duration is relatively conserved (Carvalho *et al*., 2009; Calvert *et al*., 2011; Lan *et al*., 2019; Bourdages *et al*., 2014; Khaliullin *et al*., 2018; Lee *et al*., 2018). These and other studies support the idea that ring closure arises from constriction of standardized contractile units arranged in series around the ring (Capco and Bement, 1991; Carvalho *et al*., 2009). In contrast, the kinetics and molecular requirements can be distinct among different cell types in the same animal, suggesting that multiple contractile mechanisms exist (Davies *et al*., 2018; Ozugergin *et al*., 2022; Ozugergin and Piekny, 2022). Furthermore, the kinetics of ring component abundance over the course of ring closure varies among components and across organisms: compaction and retention occur in some systems, whereas protein density remains relatively constant in other cell types (Wu and Pollard, 2005; Mendes Pinto *et al*., 2012; Khaliullin *et al*., 2018; Okada *et al*., 2021). Thus, while many aspects of actomyosin ring composition and kinetics are widely shared, some aspects of contractile mechanisms likely vary among cell types.

Various approaches to model ring closure have quantitatively recapitulated closure dynamics of cytokinetic rings in *C. elegans, S. pombe* (fission yeast), and *S. cerevisiae* (budding yeast) (reviewed in (Cortes *et al*., 2018; Pollard, 2014)). Some models utilize continuum dynamics to relate cytoskeletal protein density to the kinematics of ring closure (Zumdieck *et al*., 2007; Turlier *et al*., 2014; Sain *et al*., 2015). These approaches describe the relationship between the ring material properties and dynamics of closure without including the role of motor and crosslinker density within the ring. Further, these models lack the ability to evaluate the behavior of individual component proteins. In contrast, fully discretized physical agent-based models simulate closure of a ring made of explicitly depicted ring components, evolving those components’ position over time (Cortes *et al*., 2022; Nguyen *et al*., 2018; Stachowiak *et al*., 2014; Vavylonis *et al*., 2008). Still, these approaches do not provide a way to predict the material properties and dynamics for a particular microscopic configuration within the cytoskeletal network. A recently developed continuum mechanics model based in active gel theory relates the microscopic interactions of individual filaments to the material properties of a cytoskeletal network (Fürthauer *et al*., 2019, Fürthauer *et al*., 2021; Foster *et al*., 2022), and as such it can be used to infer microscopic network properties from macroscopic measurements.

Actomyosin rings form not only in cytokinesis but also in diverse structures within and across organisms. Contractile and stable rings are a conserved feature of germlines which, throughout phylogeny, have a complex syncytial architecture wherein many nucleus-containing compartments are interconnected via cytoplasmic bridges (Koch and King, 1969; Mahajan⍰ Miklos and Cooley, 1994; Kumar and Elkouby, 2023; Pepling and Spradling, 1998). In the *C. elegans* syncytial oogenic germline, hundreds of actomyosin rings rim the cytoplasmic bridges that connect nucleus-containing compartments to a central core of common cytoplasm, known as the rachis (Hirsh *et al*., 1976; Hall *et al*., 1999). Compartments bear such rings, called rachis bridges, throughout their lifetime. After remaining stably open for tens of hours, a rachis bridge can close if the associated compartment undergoes apoptosis, as do an estimated 50 percent of compartments (Gartner *et al*., 2008). For compartments that do not undergo apoptosis and instead become oocytes, rachis bridges close when nascent oocytes sever from the syncytium (Hall *et al*., 1999; McCarter *et al*., 1999). Interestingly, germline rings executing mitotic cytokinesis, or apoptotic or oogenic cellularization reside in a common syncytial cytoplasm and are comprised of the same conserved cytoskeletal components.

To explore the general principles of, and variations on, non-muscle contractility, we quantitatively compared actomyosin ring closure in germline stem cell mitosis, apoptosis and cellularization. We found that these different actomyosin rings within the *C. elegans* germline exhibit both distinct kinetics and distinct profiles of retention for ring proteins anillin (ANI-1) and NMMII (NMY-2). We utilized a physical framework that relates molecular scale cytoskeletal interactions to the material properties of the ring. Our model suggested that ring closure speed scales with instantaneous size rather than starting size. We also found evidence that ring closure kinetics depend on the material properties of rings, which we found to be dynamic throughout closure and unique to each subcellular context within a large syncytial cell.

## Methods

### Strain maintenance

MDX40 *C. elegans* strain in which mNeonGreen∷ANI-1 and mKate2∷NMY-2 were expressed from their endogenous loci (ani-1 (mon7[mNeonGreen^3xFlag∷ani-1])III x nmy-2(cp52[nmy-2∷mKate2+LoxP unc-119(+)LoxP])I; unc-119(ed3) III) was maintained at 20^°^ Celsius using standard methods (Rehain⍰Bell *et al*., 2017).

### Bacteria-impregnated hydrogel – Mounting Method

Worms were transferred via worm pick into a 1.5 mL tube containing 100 microliters of 0.06% levamisole in S Basal for 10 minutes. 90 - 95 microliters of 0.06% levamisole were removed, leaving the worms in 5 – 10 mL 0.06% levamisole.

Photoactivatable hydrogel solution was prepared as described (Burnett *et al*., 2018). 2-Hydroxy-4’-(2-hydroxyethoxy)-2 methylpropiophenome (Sigma Aldrich) photoactivator was suspended in S Basal (final concentrations of all components) to create a 0.001% stock (“photoactivator suspension”), and stored at room temperature, covered in aluminum foil. A 10% suspension of Poly Ethelyne Glycol (Acrylate)2 (PEGDA) (VWR, BioMatrix) was prepared in photoactivator suspension. 2 mL of OP50 liquid culture were spun at 5000 rpm for 10 minutes to pellet the bacteria. The bacteria pellet was resuspended into the photoactivator+PEGDA solution.

300 microliters of photoactivator+PEGDA+bacteria suspension was transferred first to the worm-containing tube, and then to a cell culture dish (4-chambered cover glass system #1.5 High-performance cover glass, CellVis, product number: C4-1.5H-N). Worms were pushed to the glass bottom of the dish using an eyelash tool. The dish was then carefully placed on a Speed Light Platinum Gel Documentation system, covered with aluminum foil, and exposed to UV light for 35 seconds to harden the hydrogel. 250 microliters of OP50 liquid culture were pipetted onto the cured gel containing worms.

### Fluorescence Imaging

Imaging was performed using a Nikon A1R Confocal Fluorescence Microscope (Nikon) with 1.27 NA 60x water immersion or a 1.41 NA 60x oil immersion lens and a Gallium arsenide phosphide photo-multiplier tube (GaAsP PMT) using Nikon Elements Software and Nikon Perfect Focus.

### Ring Dynamic Measurements

Actomyosin ring closure kinetics were measured by cropping the 4-D fluorescence imaging stack around the ring of interest and then performing a sum projection of the focal planes spanned by the ring. Cropped movies were registered by first manually tracking the center of the actomyosin ring using FIJI (Schindelin *et al*., 2012), and then registering the movies so that the actin ring was in the center of each frame using a custom image registration program written in MATLAB. Rings in registered movies were manually annotated using a custom script in MATLAB. A line was traced around the circumference of the ring as it closed. The pixels covered by the line and the fluorescence intensity within the pixel were recorded. The product of the line length (number of pixels) and pixel size is used as the value for circumference. The total protein was calculated as the sum of the fluorescence intensity of every pixel along the circumference of the ring. The protein density was calculated as the average of the pixels ‘fluorescence intensity.

Measurements of actomyosin ring circumference were likely underestimates of the true circumference since rings were approximated to lie within an optical section (focal plane). The true measurement is within an approximation of 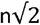 of the measured value, where n was the number of pixels that intersected the actomyosin ring.

Measurements of cytoskeletal ring circumference mNeonGreen∷ANI-1 and mCherry-NMY-2 CRISPR (LP229) [nmy-2(cp52[nmy-2∷mkate2 + LoxP unc-119(+) LoxP]) I; unc-119 (ed3) III] fluorescence intensity were collected for each actomyosin ring throughout closure for as long as accurate measurements could be made. The accuracy of actomyosin ring circumference measurements decayed as the ring completed closure in all ring types as the diminishing opening in actomyosin rings became diffraction-limited. Thus, ring circumference time series were truncated and underestimated the duration of ring closure time. Circumference time series data were smoothed by an averaging filter computed over 5 time points.

### Photo-Bleaching Correction

A photobleaching correction was performed to accurately estimate the change in fluorescence intensity of actomyosin rings over prolonged fluorescence imaging. The decrease in fluorescence intensity was measured in time-lapse image stacks using the Nikon Elements Software package. The measurements of fluorescence intensity were read into MatLab, and a lab-written script was used to correct for fluorescence intensity loss due to photobleaching. The reduction in fluorescence signal was quantified within two regions of interest (ROI); the background, and the dynamic region of interest (where actomyosin rings were measured). The average fluorescence intensity at each frame within each region of interest was tabulated. The background fluorescence intensity was subtracted away from fluorescence intensity values measured from actomyosin rings and the region of interest. An exponential function was fit to the natural logarithm transformed average intensity of the dynamic region of interest to approximate the change in fluorescence intensity due to photobleaching. The ratio method was used to correct photobleaching (Miura, 2020). The corrected fluorescence intensity value was calculated,

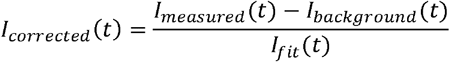

Where *I*_*corrected*_(*t*) is the bleach corrected fluorescence intensity at time t, *I*_*measured*_(*t*) is the measured fluorescence intensity at time t, *I*_*background*_(*t*) is the fluorescence intensity of the background, and *I* _*fit*_ (*t*) is the exponential fit of the fluorescence intensity in the dynamic range ROI.

### Kymograph and Montage Preparation

Kymographs were generated from registered actomyosin ring timelapse movies (see Ring Dynamic Measurement) in Image J using the PlugIn KymographBuilder (Plugins −> Kymograph −> Kymograph Builder). Montages were generated using the Fiji tool Make Kymographs (Image −> Stacks –> Make Montage).

### Population circumference and protein fluorescence intensity time alignment

Actomyosin rings within each group were aligned to the final value of each time series; that value corresponds to the last measurement of actomyosin ring circumference made before closure. The time series data for each instance of actomyosin ring closure were truncated to the duration of the shortest acquisition, so that the first time point prior to closure of the population average contained a majority of replicate measurements. The mean value of both circumference and fluorescence intensity at a given time were calculated, along with the standard error. The standard error is calculated as,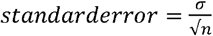, where *σ* is the standard deviation of the measurements at time *t*, relative to the last point measured, and *n* the number of measurements at time point *t* relative to the last point measured.

### Determination of the start of actomyosin ring closure

Actomyosin ring closure onset was determined by plotting the circumference time series and visually determining the time point after which the circumference curve was clearly and consistently decreasing in value.

### Average rate of actomyosin ring closure calculation

The actomyosin ring average circumference change, or closure rate, was calculated as,

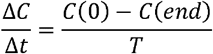

where *C*(0) is the circumference at the onset of actomyosin ring closure, *C*(*end*) is the last measured circumference, and *T* is the time between the onset of closure and the last measured value of circumference in the time series.

### Inferred time to complete closure

The inferred time to complete actomyosin ring closure, assuming that closure rate was constant, is calculated as,

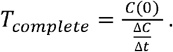

Where *C*(0) is the actomyosin ring circumference at the onset of closure, 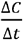 is the average rate ofclosure, and *Tcomplete* is the time in minutes to complete closure.

### Actomyosin ring protein content, and protein density time-series alignment to percentage closure

Actomyosin ring protein content and density were aligned by percentage closure. Percentage closure was calculated as,

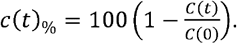

The circumferential and fluorescence intensity time series (of both ANI-1 and NMY-2) had the same length, so that the index values of elements in each measurement category corresponded to the same point in time. The sum fluorescence intensity (protein content of every actomyosin ring was then sorted into bins (groups) by percentage closure. The bins contained the associated protein content and density values of 5% increments of closure. The average and standard error of each bin, that contained protein content values organized by percentage closure, was taken.

### Actomyosin ring size and rate of ingression scaling analysis

Quantitative relationships between cytokinetic ring circumference at onset of closure and average rate of closure have been reported (Carvalho *et al*., 2009; Bourdages *et al*., 2014). This scaling law takes the form,

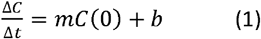

where 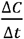 is the average change in circumference throughout closure, *C*(0) is the circumference at the onset of closure, *m* is a proportionality constant with units 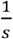, and *b* is the intercept with units of 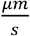. In *C. elegans* blastomeres, cytokinetic ring closure speed scales with ring size as *m* = 0.0038 *s*^−1^ and *b* = 0 (Carvalho *et al*., 2009). When not only blastomeres but also *C. elegans* vulval precursor cells are considered, this relationship is *m* = 0.0042 *s*^−1^, *b* = − 0.02 µm*s*^−1^ (Bourdages *et al*., 2014). For each cellular context of actomyosin ring closure in the germline, a range of predicted closure rates was determined using the relation from (Bourdages *et al*., 2014). The lower bound was determined using the mean value and subtracting the standard error for the value of initial circumference. The upper bound was determined using the mean value plus the standard error for the group initial circumference.

### Differential regulation of ANI-1 and NMY-2 turnover rates

We generated a parameterized model to quantitatively express the hypothesis that the turnover rate of protein on each of the actomyosin rings across cellular contexts of closure are the same. If the regulation of turnover rate is different, then the parameter values would be significantly different across cellular contexts. The hypothesis is expressed quantitatively as the change in fluorescence intensity within the rings,

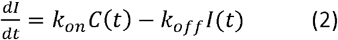

Where *k*_*on*_ is the rate of new protein that associates to the ring per unit time and length, and *k*_*off*_ is the fraction of the total protein in the ring that dissociates, per unit time. The product *k*_*on*_*C*(*t*) is the total rate of new protein incorporated into the ring, and the product *k*_*off*_*I*(*t*) is the total rate of protein lost from the ring. Integration of the rate of change in fluorescence intensity equation provides an estimate of the fluorescence intensity (total protein content) at time t,

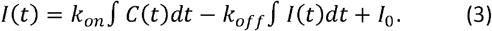

### Model Parameterization using Bayesian statistics

Bayesian analysis was used to determine whether the parameterized model (equation 3) provided an adequate description of the observed fluorescence intensity time-series (Neal, 1993; Hoffman and Gelman, 2011; Chib and Greenberg, 1995). The Markov Chain Monte Carlo method with the No U-turn sampling (NUTS) algorithm in the python package PYMC was used to generate distributions for the parameter values *k*_*on*_, *k*_*off*_, for each cellular context of actomyosin ring closure (Patil *et al*., 2010). The initial protein fluorescence intensity is a spurious parameter since the intensity time-series were normalized to the initial value. The PYMC Bayesian analysis generated the joint likelihood that parameter combinations describe the measured fluorescence intensity time-series for ANI-1 and NMY-2 given the model. The likelihoods of the parameter combinations, and the associated distributions of parameter values, provide a way of comparing the most likely parameter combinations for each context of actomyosin ring closure to one another. The model evaluated here took as input the actomyosin ring circumference and estimated the values of the parameters *k*_*on*_, *k*_*off*_, using equation (3).

The MCMC method with the NUTS algorithm was used as above to determine the distribution of parameters values for Δ*α* and *β* that described the observed ring closure curves (Figures 1, 6). The rings studied here were aligned to the last time point measured in each trajectory (see **Population circumference and protein fluorescence intensity time alignment)**. The variance among measurements was higher near the onset of closure in apoptotic and cellularizing rachis bridge closure. To account for this, we truncated the first 13 time points from the cellularization data, and the first 10 time points from the apoptosis data (as indicated along the x-axis of Figure 4A).

**Figure 1.**
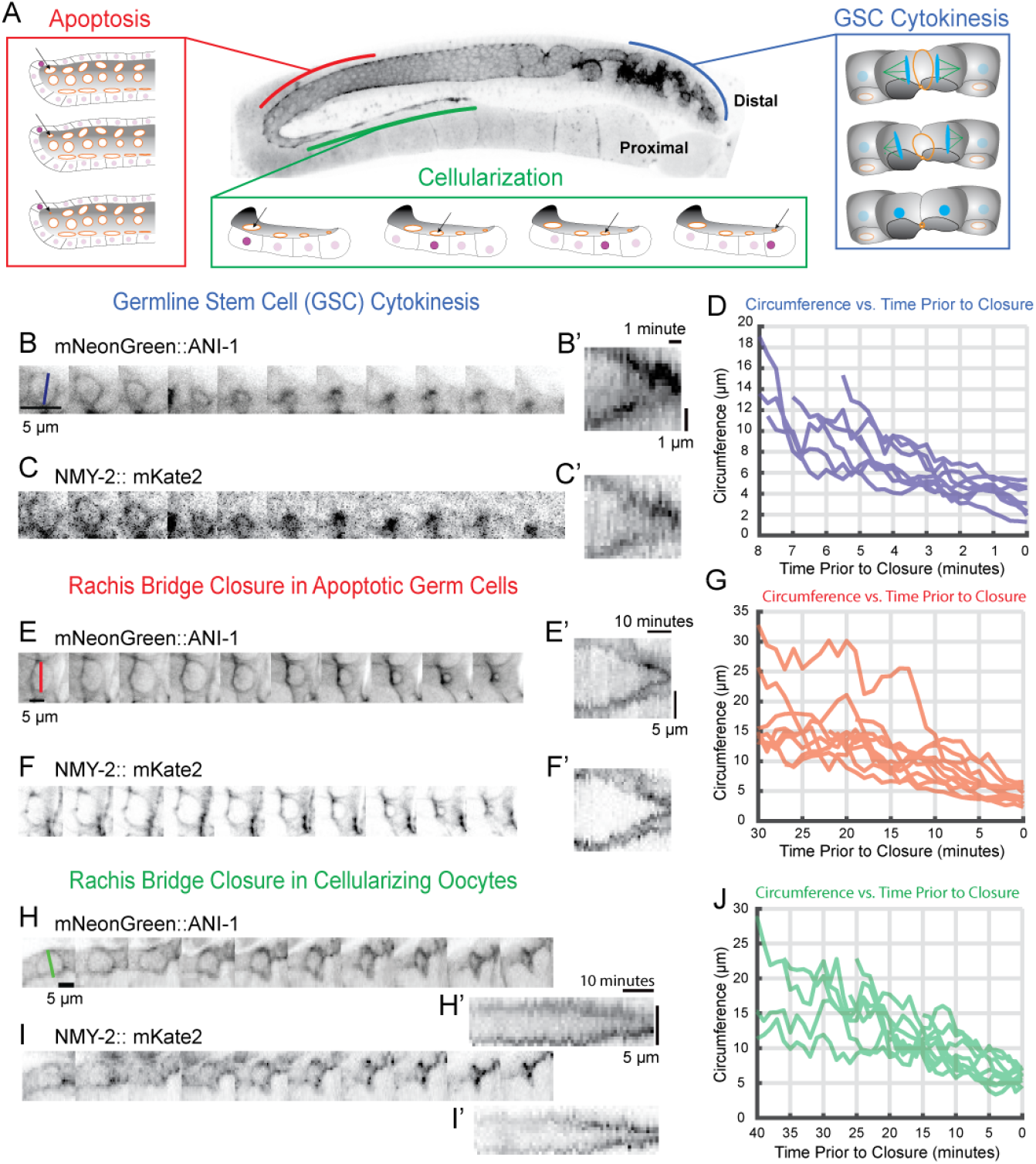
Actomyosin ring closure varies qualitatively and quantitatively among ring types in the *C. elegans* oogenic syncytial germline. A) The oogenic germline with brackets indicating the locations where germline stem cell (blue), apoptotic (red), and cellularizing nascent oocyte (green) actomyosin ring closure occur. B, E, H) ANI-1 sum fluorescence intensity image montages of actomyosin rings in each cellular context. B’, E’, H’) Kymographs of actomyosin ring closure generated from ANI-1 fluorescence sum projections. C, F, I) NMY-2 sum fluorescence intensity montages showing actomyosin ring closure in germline stem cell cytokinesis (C), apoptotic cell compartments (F), and cellularizing nascent oocytes (I). C’, F’, I’) Kymographs generated from non-muscle myosin II imaging. D, G, J) Actomyosin ring circumferential time series aligned to last recorded point. D) GSC cytokinesis (8 rings, 5 worms), G) apoptotic compartment rachis bridge closure (10 rings, 6 worms), J) cellularizing nascent oocyte rachis bridge closure (9 rings, 7 worms).

### Figures

Graphs, and boxplots were generated in MatLab. All figures were assembled in Adobe Illustrator.

### Numerical Programming

Computer programs were written in Matlab and Jupyter Notebook for Python. Source code can be found on Github under the username JackLinehan in the repository Cellular-Context-Specific-Tuning-of-Actomyosin-Ring-Contractility.

## Results

### Long-term fluorescence imaging of actomyosin ring closure in the *C. elegans* oogenic germline

Cytoskeletal rings are enrichments of F-actin, NMMII, and other structural proteins including anillin. Actomyosin rings drive a range of cellular processes in addition to canonical cytokinesis. To study contractile force generation in actomyosin rings, we leveraged the co-existence of multiple actomyosin ring closure events in a single syncytial cell: those closing in mitotic, apoptotic, and cellularizing compartments of the syncytial germline in adult *C. elegans* hermaphrodites (Supplemental Movie 1). We first studied the closure of the germline stem cell (GSC) cytokinetic ring during mitosis in the stem cell niche using fluorescently labeled conserved cytoskeletal components anillin 1 (ANI-1) (Figure 1A blue, B, B’) and NMMII heavy chain (NMY-2) (Figure 1C, C’), which enrich on the germline rachis lining, including dynamic and stable rings. GSC cytokinetic ring closure commenced as soon as enrichment of ANI-1 and NMY-2 was apparent (Figure 1B, C) (Figure 1D, Supplemental Movies 2, 3).

When germline compartments undergo apoptosis, as occurs in approximately 50% of compartments during pachytene of meiosis I prophase (Figure 1A, red) (Gumienny *et al*., 1999), the associated rachis bridge closes, separating the compartment from the common cytoplasm. Despite the importance of these rings and the knowledge that they bear many of the same components as other cytoskeletal rings, the mechanism of their closure is not known. We next used the same methods to study the closure of actomyosin rings that initially connect apoptotic compartments to the common cytoplasm (Figure 1E - G). The actomyosin rings associated with apoptotic compartments originated as anillin- and NMMII-enriched rachis bridges. After remaining at a roughly constant size for many minutes, they closed at a roughly constant rate (Figure 1G, Supplemental Movies 4-6).

In the short arm of the germline, proximal to the spermatheca, nascent oocytes cellularize from the syncytium via closure of their rachis bridge (Figure 1A, green). We also studied rachis bridge closure during the cellularization of nascent oocytes using fluorescently labeled ANI-1 (Figure 1H, H’) to track circumference (Figure 1J) and ANI-1 and NMY-2 abundance (Figure 1I, I’) (Figure 1J, Supplemental Movies 7, 8). As is the case for rings on apoptotic compartments, the rachis bridges that drive cellularization are enriched for anillin and NMMII before closure, as they remain stably open for many hours (Rehain-Bell *et al*., 2017). As rings driving cellularization closed, the rate of contraction was roughly constant (Figure 1J).

In sum, using immobilized live, intact *C. elegans* adult hermaphrodites, we observed and measured three distinct types of closing rings in the syncytial oogenic germline.

### Kinetics of germline actomyosin ring closure

Insights into the mechanisms of closure by cytoskeletal remodeling and contractile force generation can be gained from measurements of rings’ starting size, closure rate, and closure duration (Carvalho *et al*., 2009; Calvert *et al*., 2011). Towards our goal of exploring and comparing closure mechanisms of germline rings, we measured ring size over time in the three contexts (Figure 1 D, G, J). We first compared the initial circumference of rings among groups and found that GSC cytokinetic rings were the smallest (13.0 ± 3.4 µm; mean plus or minus standard deviation; Figure 2A), apoptotic germ cell compartment actomyosin rings were approximately 1.3 times larger than GSC cytokinetic rings (17 ± 5.0 µm; Figure 2A), and cellularizing germ cell compartment actomyosin rings were roughly 1.6 times larger than GSC cytokinetic rings (21 ± 5.50 µm; Figure 2A).

**Figure 2.**
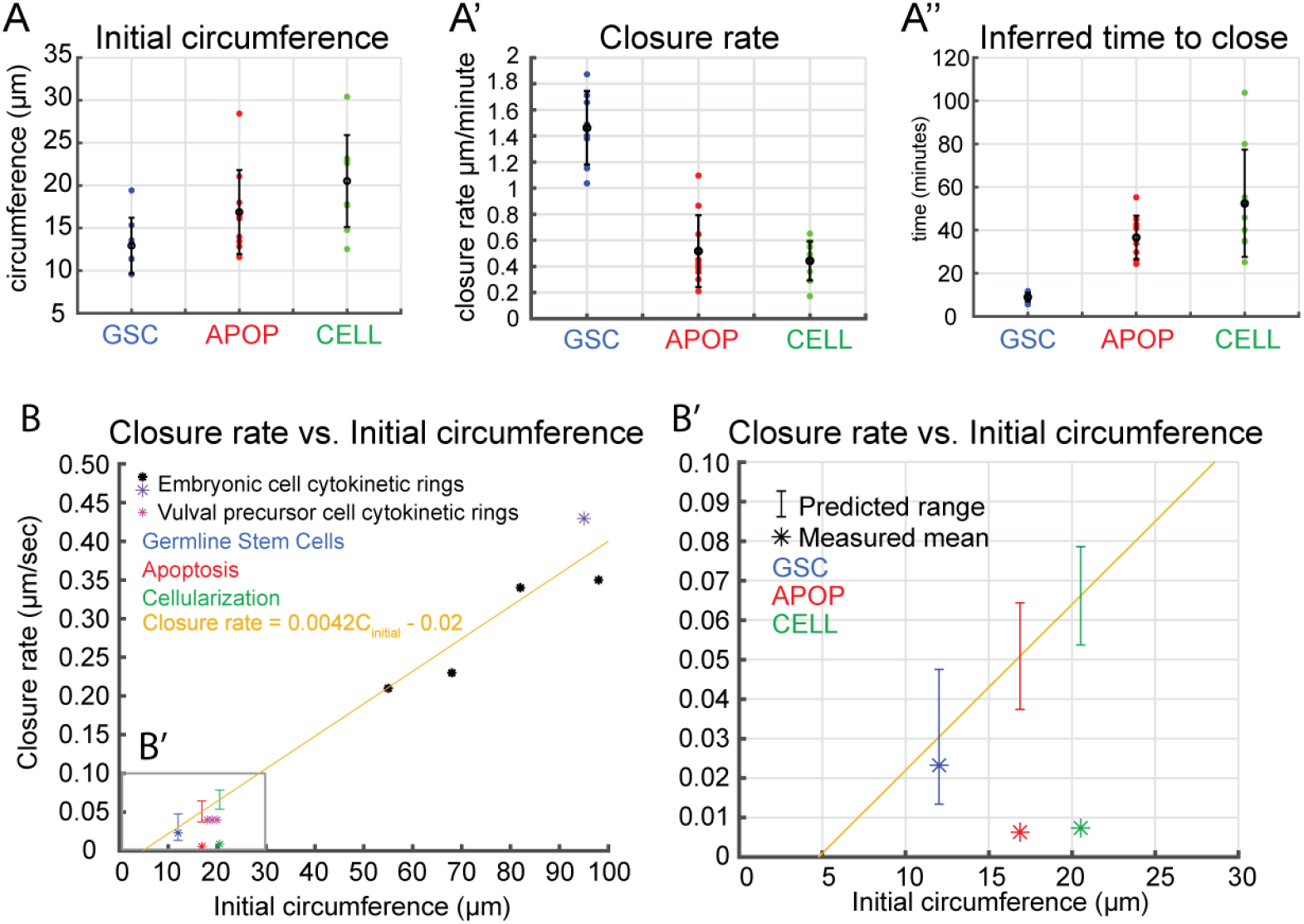
Comparison of actomyosin ring closure in mitotic germline stem cells, apoptotic, and cellularizing germ cell compartments. A) Actomyosin ring circumference at the onset of closure (mean ± standard deviation in black). A’) Actomyosin ring closure rate (mean ± standard deviation in black). A’’) The inferred time to close (mean ± standard deviation in black). B) Starting ring size and closure rate for multiple *C. elegans* cell types. Closure rates for embryonic stage cytokinesis and cytokinesis in vulval precursor cells were derived from the literature (Carvalho *et al*., 2009; Bourdages *et al*., 2014) B’) Predictions of actomyosin ring closure rates based on actomyosin ring scaling relationship; brackets: range of expected ring closure speed; blue, GSC cytokinetic rings; red, apoptosis; green, cellularization.

We next compared closure rates and found that the average closure rate was the fastest, at 1.46 ± 0.28 µm/minute, in GSC cytokinetic rings (Figure 2A’). The apoptotic actomyosin ring closed at an average of 0.52 ± 0.27 µm/minute (Figure 2A’) and cellularizing rings displayed an average closure rate of 0.44 ± 0.15 µm/minute (Figure 2A’). Since closure rate was roughly constant for all ring types, we inferred duration of ring closure by dividing the initial circumference by the average ring closure rate. The average inferred time to close in GSC cytokinetic rings was 9.0 ± 2 minutes (Figure 2A’’). The average time for the actomyosin ring to close in apoptotic germ cell compartments was 37.0 ± 10 minutes (Figure 2A’’). The average inferred time to close in cellularizing actomyosin rings was 52.0 ± 25 minutes (Figure 2A’’). Our findings clearly demonstrate that the three ring types in the germline exhibit different speed and closure times.

Cytokinetic ring kinematics have previously been explained by a scaling law relating the initial circumference to closure rate (Carvalho *et al*., 2009; Calvert *et al*., 2011; Bourdages *et al*., 2014). We explored whether starting ring size was sufficient to predict closure speed for the three *C. elegans* germline ring types (Figure 2 B, B’)(see Materials and Methods). The measured rate of GSC cytokinetic ring closure indeed fell within the range of closure speed predicted by starting size. Conversely, apoptotic or cellularizing rachis bridges closed with substantially lower than rates previously predicted from relating initial ring circumference to rate of closure (in other *C. elegans* cell types) (Figure 2B’). We conclude that of the three ring types investigated here, the kinetics of only GSCs are compatible with prior theories.

### Force balance and kinetics of cytoskeletal rings

To understand the physical differences among ring types that led to their divergent kinetics (Figure 2), we next tested whether a simple physical model for ring closure could recapitulate our measurements when distinct sets of fitted parameter values were implemented (see Figure 4). We modeled the ring as a viscous active material, whose motion is generated by the competition between passive viscous stresses and active stresses generated by motor molecules and crosslinkers. In our model the network stress takes the form

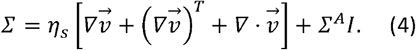

where *η* _*s*_ denotes the network viscosity, *Σ*^*A*^ is the scalar active stress and *I* is the identity matrix (see Table 1). The force balance equation of this structure is *∇· Σ*=0. Depicting the ring as a thin ribbon of active gel (see Figure 3A), and imposing azimuthal symmetry, this leads to

**Table 1.**
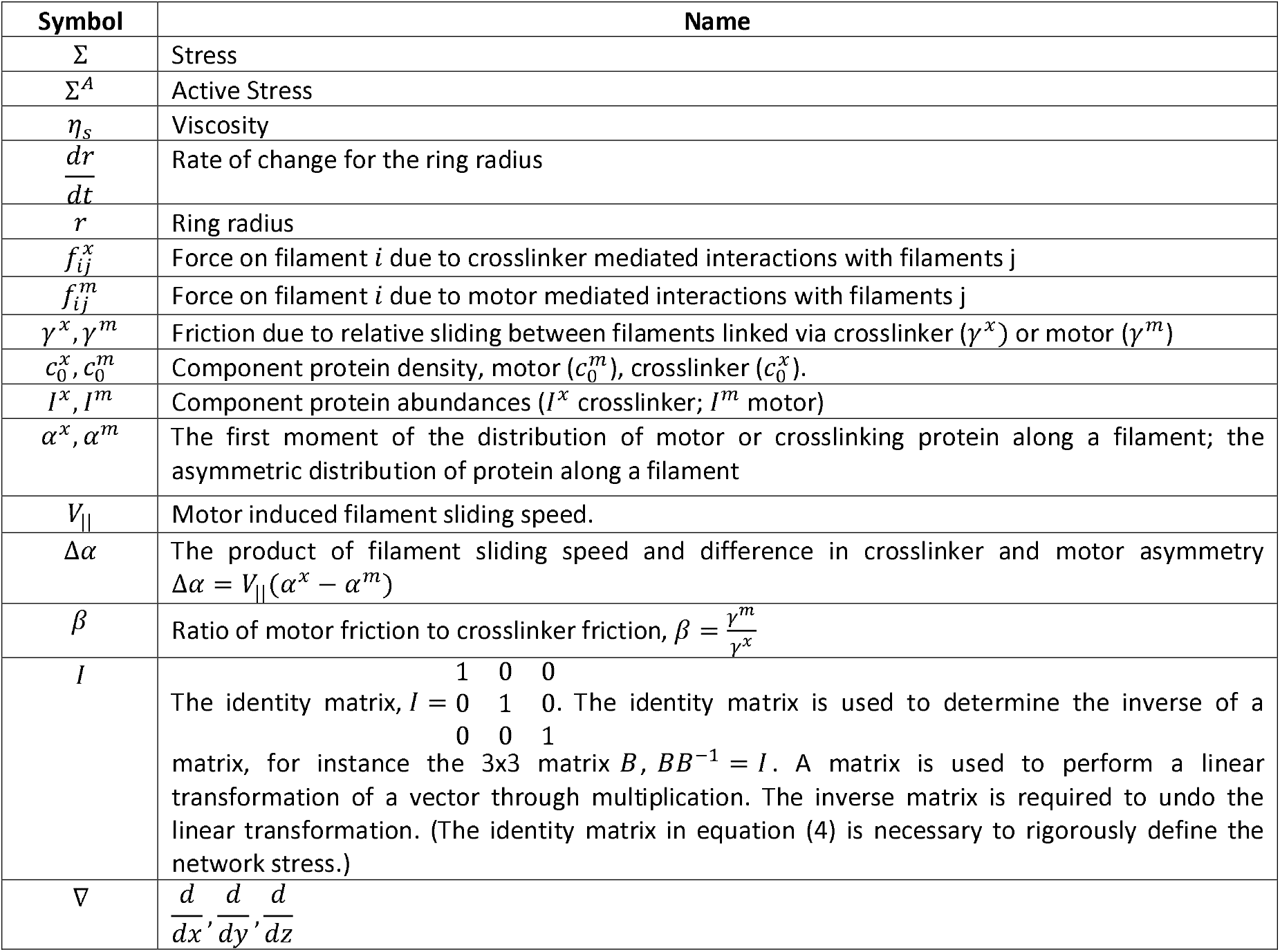

**Figure 3.**
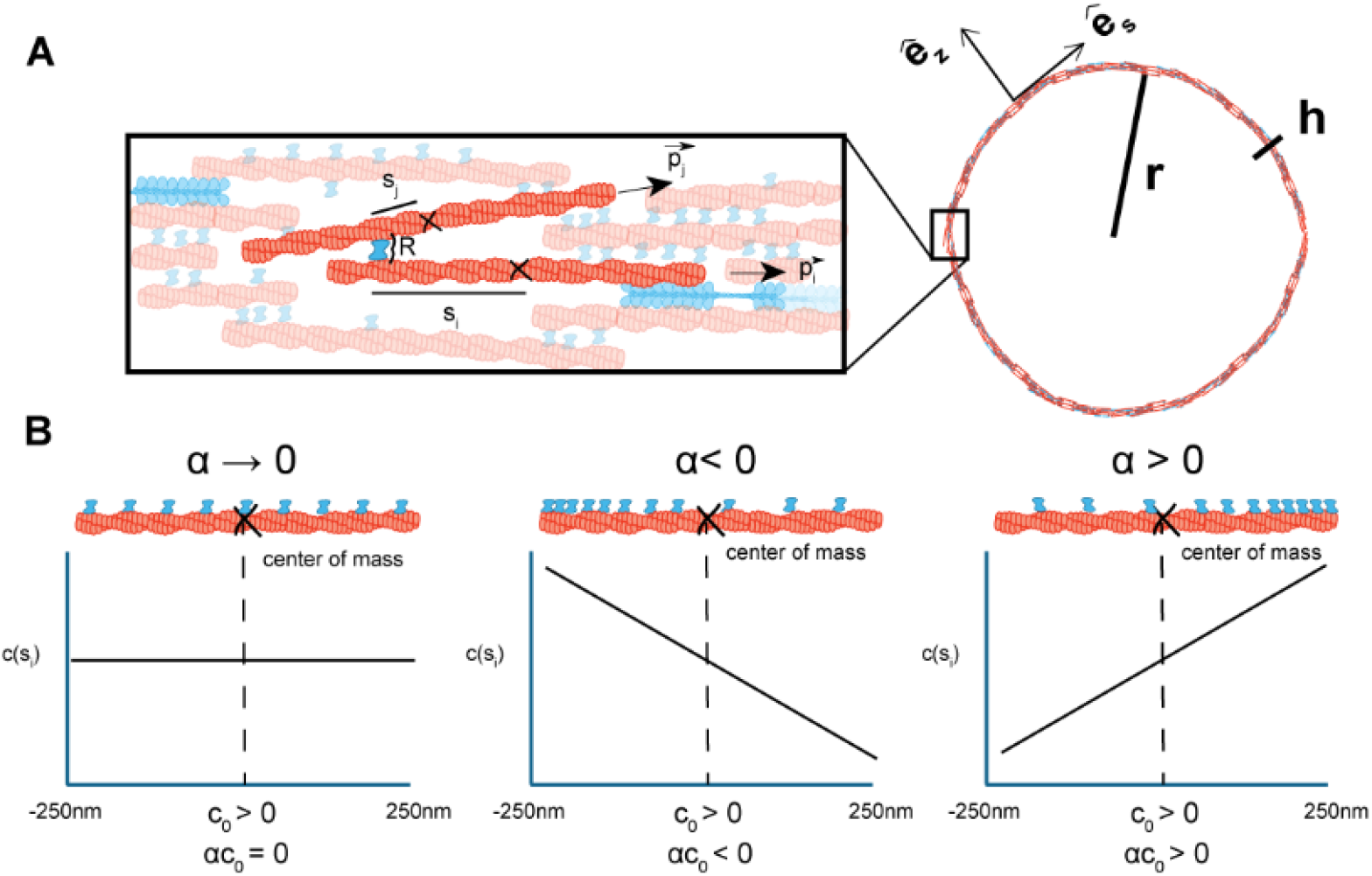
Theory for actomyosin rings. A) An actomyosin ring with radius *r* and width *h* sis described with the normal unit vector 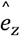 and tangent unit vector 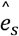. Schematic of forces exerted on filament *j* by neighbor filaments *I* through motor or crosslinking head of width 13 nm. The force experienced by filament *i* depends on the distance of the motor or crosslinker head from the respective filament’s center of mass. Each filament has an orientation described by the unit vector 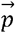. B) Schematic of protein distribution along F-actin length when the distribution of protein is uniform, and asymmetrically distributed protein along F-actin, and corresponding *α*.

**Figure 4.**
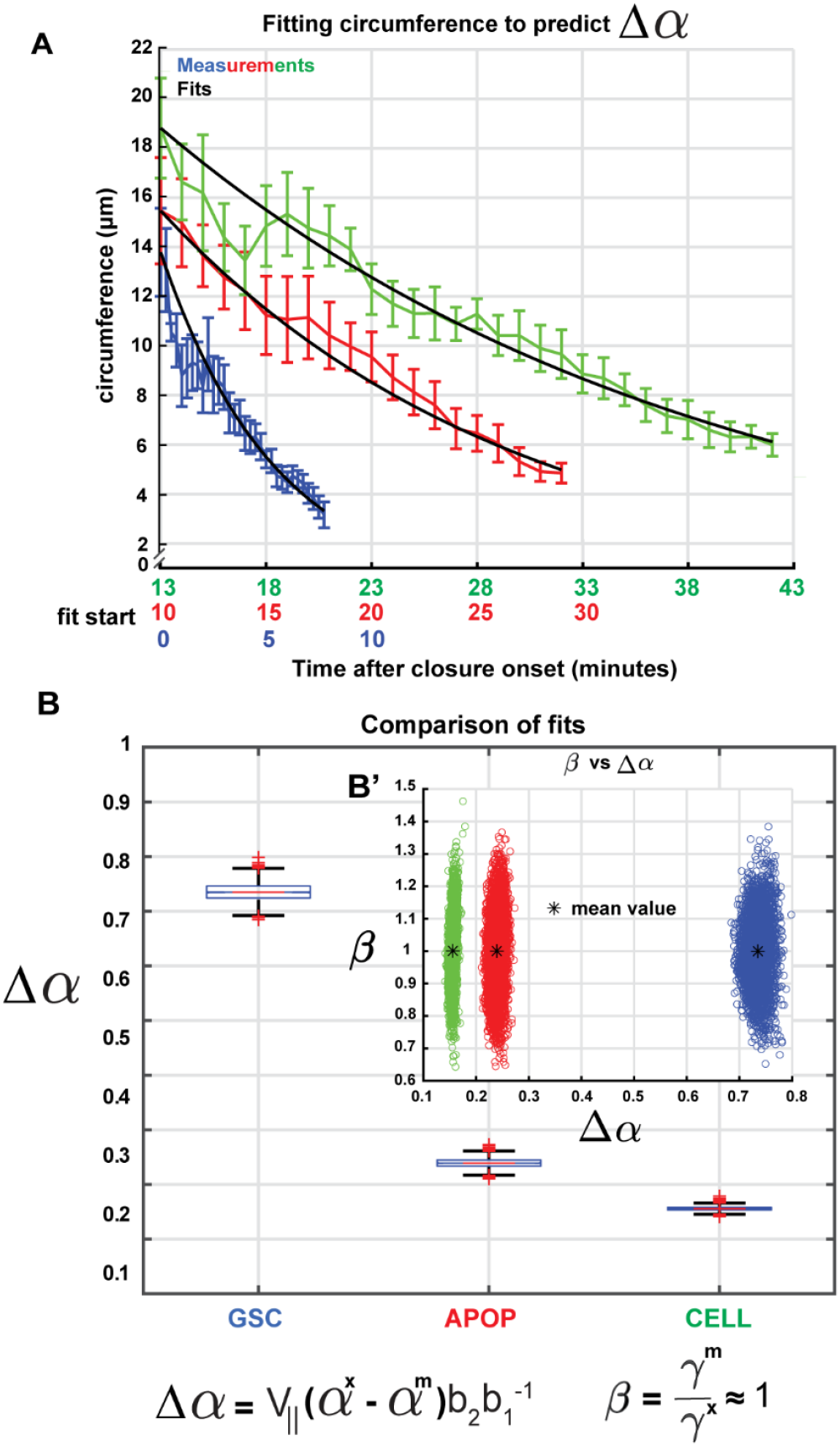
Minimal model and the evolution of ring contractility. A: Comparison of measured circumferences in time (Blue, GSC Cytokinesis; red, apoptosis; green, cellularization; mean ± standard error) with predicted circumference generated by parameterized model (black). B) Boxplots of Δ*α*value distributions determined by MCMC method. The central line (red) of the boxplot indicates the median of the distribution. The top side of the box indicates the 75^th^ percentile, and the bottom side indicates the 25^th^ percentile. The whiskers indicate the maximum and minimum values of the distributions, and the red crosses indicate outliers. B’) Posterior distributions of *β* and Δ*α* generated using the Markov Chain sMonte Carlo Method.

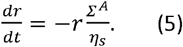

Details of the derivation are given in the Appendix. Importantly, according to this equation, the current rate of contraction of the ring is proportional to its current size, and to the ratio of its active stress to its viscosity. Thus, to interpret our results of ring size as a function of time in the context of our physical model, we next related the active stress and viscosity of rings to their molecular composition.

### Linking molecular and material properties of cytoskeletal rings

To explore the molecular underpinnings of ring contractility 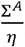, we deployed a theoretical physical framework that relates molecular scale interactions to filament network material properties (Fürthauer *et al*., 2021) (Fürthauer *et al*., 2019). The theory derives material properties from component abundance, beginning with a description of the effective forces acting on an actin-like filament *i* due to interactions with neighboring filament *j* via connections made by crosslinkers representing non-motor crosslinkers including anillin, and motors representing NMMII (Figure 3B).

For a number of passive crosslinkers *c*^*x*^(*s*_*i*_,*s*_*j*_), bound between points *s*_*i*_ on filament *i* and point *sj* on filament *j*, the effective force 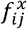 exerted between the filaments is assumed to be dominated by friction. In other words, the effective force is proportional to the velocity difference between attachment points *s*_*i*_ and *s*_*i*_

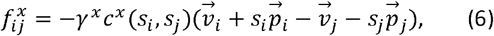

where *γ*^*x*^ is the friction constant for crosslinkers, 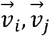 describe the velocity of the respective filaments and 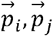 their orientation (Figure 3A). In the case of a motor abundance *c*^*m*^(*s*_*i*_,*s*_*j*_), in addition to the frictional coupling of filaments, we expect that the presence of active forces relates to motor stepping. Thus, the force 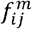 exerted between filament *i* and *j* in the simplest case of a linear force-velocity curve for the motor, will read

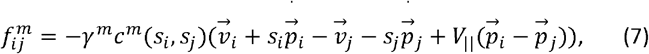

With *γ*^*m*^ being the friction constant for motors and *V*_‖_ being the motor stepping speed.

To allow for a non-uniform distribution of crosslinkers and motors, we postulate that their intensities differ linearly along the filament length *L* as follows

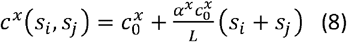

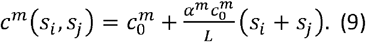

In the above expressions, the *s*_*i*_ and *sj* parametrize the positions on the filament and have values between − *L*/2 and *L*/2. 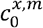 is the total amount of crosslinker or motor per filament. The parameters *α*^*x,m*^quantify the distribution of the crosslinker and motor proteins along the filament length, respectively. Positive values of *α*^*x*^ and *α*^*m*^ indicate that the crosslinkers or motors cluster near the end that the motor walks towards, while negative values indicate the opposite.

Following an established coarse graining procedure (Fürthauer *et al*., 2021), we obtained expressions for the network viscosity *η*_*s*_ and the network active stress *Σ*^*A*^, which reads

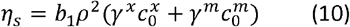

And

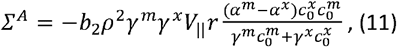

respectively. Here *ρ* is the density of actin filaments. The constants *b*_1_ and *b*_2_ depend on relative dimensions of the ring components (see Appendix). Together, equations (8, 9, 10, and 11) lead to recasting the ring dynamic equations of motion as

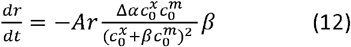

where Δ *α*=*V*_‖_(*α*^*x*^− *α*^*m*^),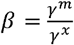and 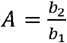.

Equation (12) relates ring contraction behavior to two effective fit parameters (Δ*α,β*) and protein abundances 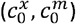 on the ring. Δ*α* and *β* reflect microscopic interactions within the filament network that cannot be directly measured (see Figure 4). The total abundance of the crosslinker 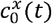 and the motor 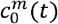 at time *t* can be linked to the measured total fluorescence intensities of ANI-1, *Ix*(*t*), and NMY-2, *Im*(*t*) (see Figure 5A-C’), via the expression

**Figure 5.**
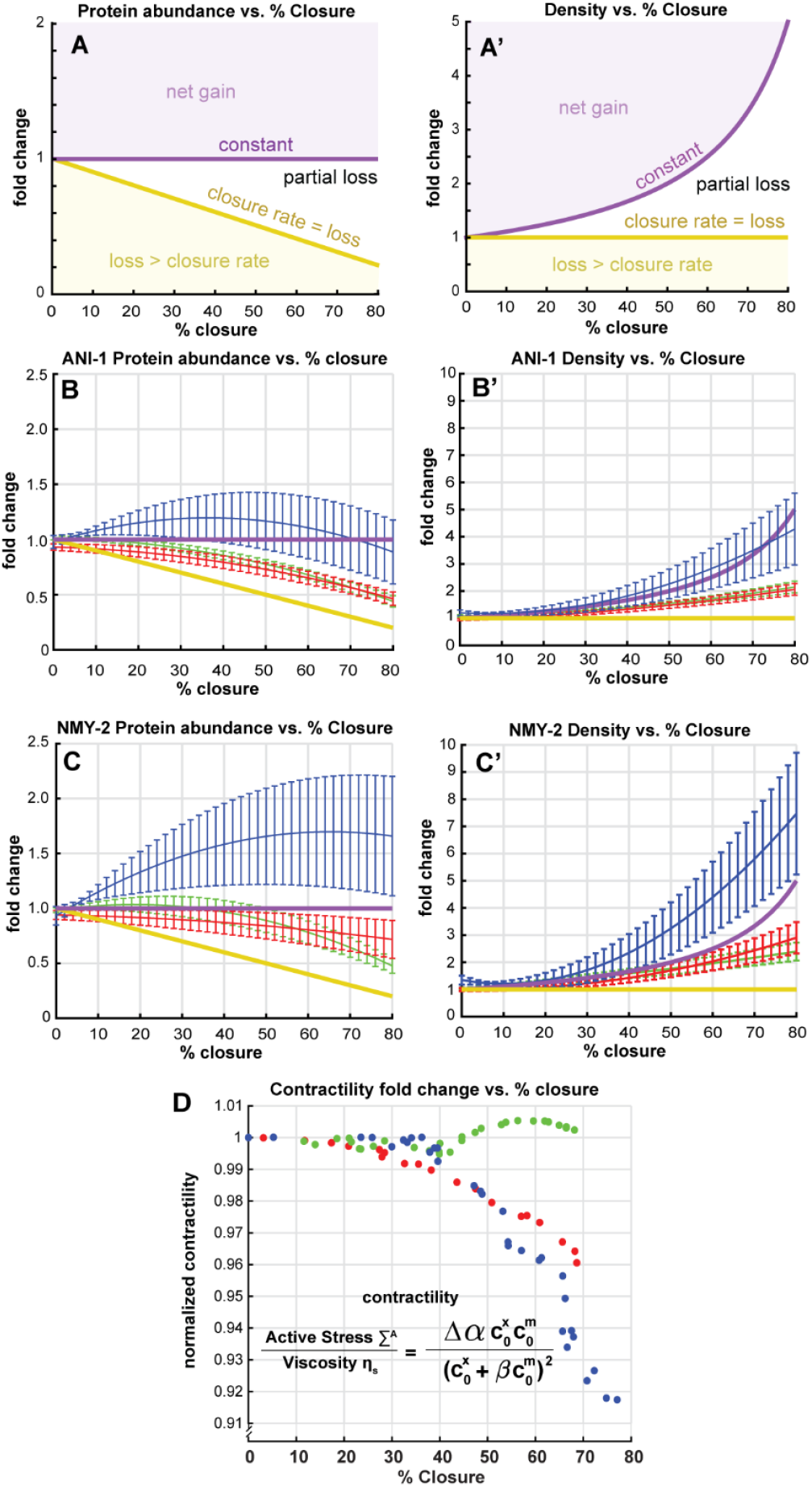
ANI-1 and NMY-2 density and abundance during closure. A, A’) Limiting cases of protein turnover; A’: expected fold change in protein abundance as the ring closes given a net gain in protein (light purple), constant abundance (purple), fractional loss (white), equivalency between protein change and circumference change (yellow), and when protein loss is greater than rate of closure (light yellow). ANI-1 abundance as a function of percent closure; GSC cytokinesis (blue), apoptosis (red), cellularization (green)(mean ± standard error). B’) ANI-1 density as a function of percent closure (mean ± standard error). C) NMY-2 abundance as a function of percent closure (mean ± standard error). C’) NMY-2 density as a function of percent closure(mean ± standard error). D) Fold change in actomyosin ring contractility throughout closure; GSC cytokinesis (blue), apoptosis (red), cellularization (green).

**Figure 6.**
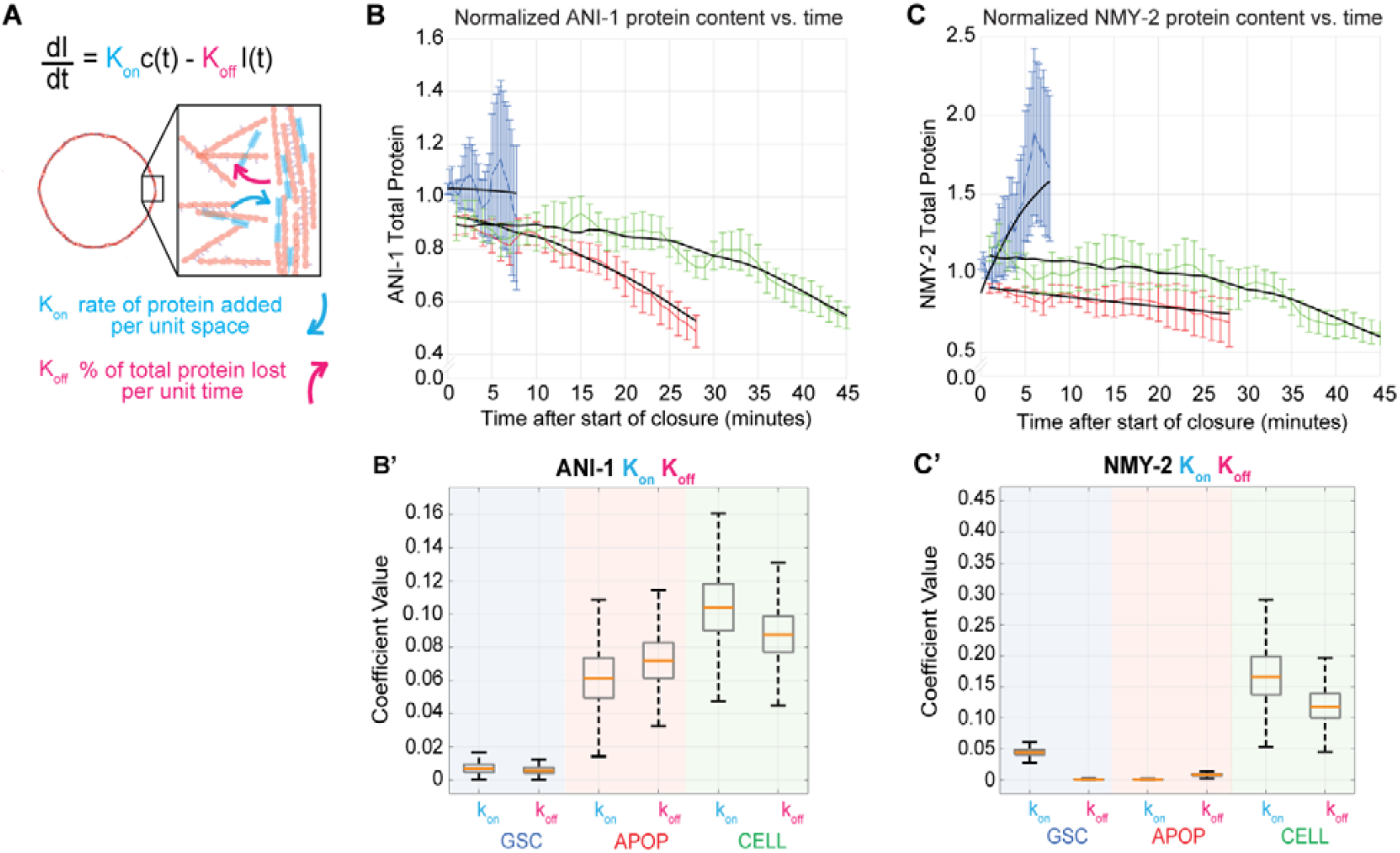
Time-dependent model of ANI-1 and NMY-2 abundance throughout ring closure. Protein content estimates are normalized to initial fluorescence intensity. The mean value of protein fluorescence intensity at each time and standard error is shown. A) Model schematic: the change in the fluorescence intensity, 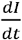, in pixels around the circumference of the ring is due to changes in the number of fluorescently labeled molecules within the area of a pixel. In the model, the change in the fluorescence intensity is dependent on a rate of protein flux onto the ring per unit area per unit time *k*_*on*_, and a parameter *k*_*off*_ describing the fraction of the total protein shed from the ring per unit of stime. B) GSC cytokinesis (blue), apoptotic cell compartment (red), cellularizing nascent oocyte (green) (mean ± standard error); ANI-1 normalized protein fluorescence intensity population time series, black lines: model prediction generated using the mean value of the distribution of likely values for *k*_*on*_ *k*_*off*_. NMY-2 normalized protein fluorescence intensity population time-series, GSC (blue), apoptosis (red), cellularization (green) (mean ± standard error), with model predictions generated from the mean value of likely parameter values for *k*_*on*_,*k*_*off*_ (black). B’, C’) Coefficient value distributions for *k*_*on*_,*k*_*off*_ determined using the MCMC method. Boxplots in (B’, C’) have an orange line indicating the median, the top of the box is the 75^th^ percentile, the bottom of the box is the 25^th^ percentile, the whiskers indicate the max and min values of the distribution.

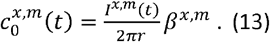

Here we introduced the conversion factors *β*^*x,m*^, that relate fluorescence intensity to protein amount.

With this, equation (12) can be recast as

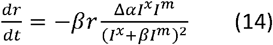

Note Δ *α*=*AV*_‖_(*α*^*x*^− *α*^*m*^)and 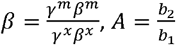. Note that the form of equation (14) is identical to that of equation (12). The additional parameters introduced to convert fluorescence intensity to protein amount have been incorporated into the two fit parameters Δ*α*, and *β*.

Equation (14) relates measured changes in fluorescence intensity of crosslinker and motor abundance to the rate of change of the ring radius. The model has two effective free parameters (Δ*α*, and *β*) that must be obtained by fitting to measurements. Doing so (see Figure 4 below) is predicted to provide insights into the underlying cytoskeletal interactions that result in force generation during Sring closure.

### Differences in the asymmetric distribution of cytoskeletal components along actin-like filaments is sufficient to recapitulate observed ring closure trajectories

The active gel theoretical framework, resulting in the coupled system of equations (2) and (14), allowed us to explore what microscopic details of network architecture determine the kinetics of and motors is equal (*γ*^*x*^ ≈ *γ*^*m*^ such that *β* ≈ 1). This assumption was substantiated by demonstrations actomyosin ring closure. We introduced a simplification, that the friction (with F-actin) of crosslinkers that myosin motors under load interact with F-actin via catch-slip bonds, such that NMMII bundles Stam *et al*., 2015). We also assumed that the motor or crosslinker asymmetry parameter Δ*α*, is constant behave similarly to crosslinkers (Cortes *et al*., 2020; Coluccio and Geeves, 1999; Erdmann *et al*., 2013; in time and only the overall protein abundance *I*^*x,m*^(*t*)changes. We employed the MCMC method to determine the distribution of values of the parameter Δ*α* that relates the measured protein density Δ*α* = 0.73 ± 0.07 allowed a good fit to the observed ring closure kinetics (Figure 4A, B-B’ black: model dynamics (see Figure 5A-C’) to the observed ring closure curves. For germline stem cell cytokinesis, predicted curves using the mean Δ*α* value). For apoptotic or cellularizing ring closure, Δ*α* = 0.24± 0.03, and Δ*α* = 0.16 ± 0.004 (mean ± std), respectively (Figure 4A, B-B’). Additionally, the mean values of the *β* distributions were indistinguishable for the three ring types (approximately 1; Figure 4B’). Thus, the MCMC method revealed that measured ring kinetics could be fitted well with unique Δ*α* values for each of the three ring types. Thus, while the structural components of all three ring types are shared, what may vary among contexts is the distribution of actin binding proteins along F-actin, a factor possibly tuned by cytoskeletal regulators. Importantly, our model suggests that divergent ring closure kinetics can result from the differential recruitment and/or retention of ring component proteins (see Figure 6).

### ANI-1 and NMY-2 density evolution distinguishes the GSC cytokinetic ring from rachis bridges

Defining ring speed in these terms (Eq. 14) predicts that differences in ring closure speed among ring types reflect differences in protein abundance dynamics. Specifically, since GSC cytokinetic rings are significantly faster than rachis bridges in apoptosis or cellularization, the time-evolution of protein abundance is predicted to be different between GSC rings and the other two classes of rings. To test these predictions, we measured protein abundances of the motor protein NMY-2 and crosslinker ANI-1 in rings throughout contraction. We interpreted our protein abundance measurements by comparing them to calculated density change for various theoretical regimes of recruitment, retention, and loss (Figure 5 A, A’). If the net amount of protein is constant (if all protein is retained) throughout closure, 80% closure results in a 5-fold increase in density (Figure 5 A, A’; light purple). If protein is lost at a rate that is exactly equal to the rate of closure, then the density would remain constant (Figure 5 A, A’; purple line). A change in density greater than five-fold indicates a net gain in the ring; a change between one- and five-fold corresponds to a partial loss of protein.

Normalized fluorescence density and abundance of tagged ANI-1 and NMY-2 varied among the three ring types. GSC cytokinetic rings experienced a greater fold change in density of ANI-1 and NMY-2 than the two types of rachis bridges, which exhibited similar fold changes (Figure 5B’, C’). ANI-1 abundance initially increased during GSC cytokinesis and began to decrease around 50% closure. In both types of rachis bridges, ANI-1 was lost throughout closure (Figure 5B). NMY-2 abundance increased in GSC cytokinetic rings and decreased in both instances of rachis bridge closure (Figure 5C). These measurements support the idea that ring dynamics are determined by ring type; differences in contractile regulation may underlie kinetic differences.

### Actomyosin ring contractility exhibits unique dynamics across subcellular contexts

Our physics-based framework revealed that the closure speed of a ring at any time is determined by two distinct quantities (see equation (11)): the ring’s radius *r* and its material properties that are summarized in the contractility ratio 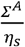. As discussed above, both the active stress *Σ*^*A*^ and *η*_*s*_ viscosity can be dynamic, affecting the overall kinematics of the ring closure. To estimate each ring type’s contractility as a function of time, we used the fitted values of Δ*α*, which varied among ring types (Figure 4B), and *β*, which did not (Figure 4B’). We calculated contractility for each ring type from equation (14),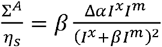 using the mean value of the fit distributions of Δ*α* and *β* (Figure 4 B-B’).

We found that the GSC cytokinetic ring experienced a ~10% reduction in contractility during closure (Figure 5D; blue). Apoptotic compartment rings experienced a similar, but more gradual, reduction in contractility (Figure 5D; red). By contrast, in cellularization rings, contractility remained constant throughout closure (Figure 5D; green). These results suggest that the material properties of actomyosin rings are dynamic throughout closure and diverge among cellular contexts. These findings further support the conclusion that the underlying mechanics of ring closure are conserved, and that the magnitude of ring contractility can be tuned via biochemical regulation of component protein abundance and distribution within the filament network.

### Time dependent ANI-1 and NMY-2 abundance dynamics differ among subcellular contexts of actomyosin ring closure

Above, we characterized ring state as a function of percent closure (normalizing instantaneous ring size to starting size) to consider composition dynamics at similar extents of network contraction. By contrast, characterizing ring dynamics as a function of time enables quantitative analysis of the temporal intermediates of contractility. Since the time to complete ring closure varied several-fold across subcellular contexts we next asked how protein abundance varied as a function of time. The total amount of the scaffold protein ANI-1 remained relatively constant in the GSC ring as it quickly closed (Figure 5B, C, blue). During the slower closure of apoptotic and cellularizing rings, total ANI-1 gradually diminished (Figure 5B, C, red: apoptosis, green: cellularization). Total abundance of the NMMII heavy chain NMY-2 increased in the GSC ring during closure, stayed relatively constant during apoptotic ring closure, and gradually decreased in cellularization ring closure (Figure 5B, C).

We next leveraged our time-resolved abundance measurements to estimate turnover rates for each protein in each ring type. We used a generalized model for the change in protein content over time:

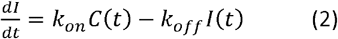

where material incorporation was the product of *k*_*on*_ (the rate of protein added per unit space per unit time) and the current circumference of the ring *C*(*t*) and material loss was the product of *k*_*off*_ (the fraction of ring protein shed per unit of time) and the current protein content, *I*(*t*) (Figure 6A). We calculated the likelihood that the measured fluorescence intensity time-series data were generated by the linear model given a normally distributed error. The Markov Chain Monte Carlo (MCMC) method was used to determine the distribution of parameters *k*_*on*_ and *k*_*off*_ that fit each average fluorescence intensity over time curve. As expected, the predicted fluorescence intensity time series calculated from the mean parameter values for *k*_*on*_ and *k*_*off*_ (Figure 6B, C, black lines) agreed well with our measured fluorescence intensity time series. The distributions of the parameter values *k*_*on*_ and *k*_*off*_ for ANI-1 and NMY-2 were well separated among the three ring types (Figure 6B’, C’). Specifically, ANI-1 *k*_*on*_ and *k*_*off*_ were very different in germline stem cell mitosis than in either rachis bridge type, and *k*_*on*_ and *k*_*off*_ of NMY-2 were all different among the three ring types (stat test results?). This suggested that the regulation of both ANI-1 and NMY-2 differs among the three ring types studied here. This finding provided further support for the idea that a unified model framework can explain different rings’ behavior, given ring-specific evolutions of component protein abundances in time (Equation 14).

## Discussion

We sought unifying principles for how contractility emerges from the cytoskeletal interactions within non-muscle actomyosin networks. To do so, we compared three instances of actomyosin ring closure within a large syncytial cell. We observed actomyosin rings in germline stem cell cytokinesis, the closure of apoptotic compartments, and the cellularization of nascent oocytes (Figure 1). We found that the kinetics of ring closure varied among the three groups and that only GSC cytokinesis could be described using prior theories (Figure 2). We used a theoretical framework to model germline ring closure and found evidence that rings in diverse settings close via a shared mechanism and that kinematic differences result from context-specific regulation of ring component turnover and mesoscale network architecture.

### Variation in the magnitude of contractile stress helps explain the varied kinematics of ring types

A scaling “horsepower” relationship has been proposed to explain how, across a range of cell types, the speed of ring closure scales with starting size: The starting circumference is thought to dictate the starting abundance of ring material, such as if ring components were arranged in a series of contractile units, and the number of units acting in series dictates speed (Capco and Bement, 1991; Carvalho *et al*., 2009; Calvert *et al*., 2011; Mendes Pinto *et al*., 2012; Guillot and Lecuit, 2013; Cuvelier *et al*., 2023). This relationship is expressed as 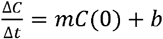 (equation 1; see Materials and Methods), where the proportionality constant *m* is determined via fitting (Carvalho *et al*., 2009; Calvert *et al*., 2011; Bourdages *et al*., 2014). Here, we found that the rate of ring closure in GSC cytokinesis was accurately predicted using the size-speed scaling relationship with both embryonic and larval somatic C. elegans cell types (Bourdages *et al*., 2014), indicating that the mechanics of cytokinesis are broadly conserved across cell type and size at least within this species.

The same relationship failed to predict the observed closure rates of the two types of rachis bridges (Figure 2). Those data, in combination with our measurements of fluorescence density and abundance within germline rings (Figures 5, 6), suggested that the proportionality constant *m* contains information relevant to ring type (i.e. cytokinetic rings). Fitting ring kinematics of all three types of germline rings using our active gel-based theory revealed that the proportionality constant *m* captures the material properties of the ring (Figure 4, 5D). Indeed, by using equation (15) with an asymmetry (finite Δ*α*), we can relate the constant *m* to the microscopic properties of the network as follows (for details see the appendix)

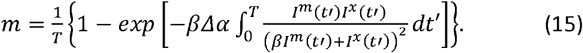

Thus, in addition to the protein kinetics affecting the fluorescence intensities *I*^*m,x*^(*t*), the values of the product Δ*α* also affect ring closure kinematics. Furthermore, they do so in an exponential manner, highlighting their prominent role. This confirms our intuition that the proportionality constant *m* contains information about the closure mechanism; that contractility is determined by ring components’ abundance and distribution along filaments. In sum, the dimensions of a contractile array captured by the initial circumference *C*(0) are important to predict closure kinematics, in keeping with the scontractile unit or “horsepower” hypothesis, but material properties, which vary not only among ring types but over time within a ring type (Figure 4C), are also necessary for explaining contractile speed.

### The asymmetric distribution of NMMII along F-actin and its motoring tune ring contractility

We generated a physical model to test how our measurements of ring kinetics relate to mechanical differences among ring types (Figure 3). Our model for ring closure rate (Equation 5) states that contraction rate is proportional to current ring size and ratio of active stress to viscosity. We related the ring active stress and viscosity to ring molecular composition (Equations 5-14) and found that the ring closure rate depended on two fit parameters (Δ*α,β*) and the densities of the crosslinking and motor proteins 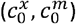 (Equation 12).

Non-motor crosslinkers including anillin can generate contractile forces in reconstituted F-actin networks but only when the level of filament overlap is low (Kučera *et al*., 2021); in cytokinetic rings Factin is thought to be maximally overlapped (Swulius *et al*., 2018; Mangione and Gould, 2019). Therefore, the non-motor crosslinker may contribute minimally to contractility. *In vivo* and in our model, NMMII generates contractile stresses via motoring and contributes to network viscosity by crosslinking F-actin (Wang *et al*., 2020; Maddox *et al*., 2007; Ding *et al*., 2017). The contribution of NMMII motoring to ring closure *in vivo* varies across organismal contexts (Lord *et al*., 2005; Osório *et al*., 2019; Wloka and Bi, 2012) and could vary among the three subcellular contexts we explored. Since the magnitude of the force generated by NMMII depends on both its polarized distribution along filaments and its motor activity (Equation 5), either the asymmetric distribution, degree of filament sliding, or a combination of both, could contribute to tuning the contractility of rings across cellular contexts.

### The asymmetric distribution of crosslinking and motor protein along F-actin is a physical inevitability

Our work indicates that the generation of contractile stresses depends on the asymmetric distribution of crosslinking and motor protein along the length of actin filaments. The parameter *β*was very similar across the three rings and indicates that friction and relative fluorescent intensity per motor and crosslinker are not substantially different among ring types. Given the conserved behaviors of NMMII and ANI-1 across cellular contexts, and that we fit distinct values of Δ*α* (Figure 4) to each ring type, we propose that Δ*α* captures features of the rings relevant to the biochemical regulation of contractility in each cellular context. The interactions between cytoskeletal proteins are known to directly impact network architecture (reviewed in (Kadzik *et al*., 2020)). Asymmetry likely stems from the motility of NMMII to actin barbed ends and the barbed-end-dwelling of non-motors including anillin (Kučera *et al*., 2021)). Furthermore, mutually exclusive zones of actin populated by NMMII and non-motor crosslinkers may arise, given that NMMII forms bundles with more widely-spaced actin filaments than those created by many non-motor crosslinking proteins (Sinard *et al*., 1989). The crosslinking proteins fascin and *α*-actinin segregate due to differences in size, occupying mutually exclusive binding zones along filaments within bundles (Winkelman *et al*., 2016; Li *et al*., 2015). Formation of such sorted zones of NMMII and non-motor crosslinkers along actin filaments within actomyosin rings could result in network architectures that limit drag on filament sliding. The polarization of motor and non-motor crosslinkers along F-actin likely also depends on filament length, age, and nucleotide content; F-actin polymerization dynamics could thus tune ring contractility by modulating crosslinker/motor sorting. Depletion of F-actin regulators including cofilin, formin, and capping protein may provide insight into the role of F-actin length in determining the polarized distribution of component proteins within rings. Our estimated turnover rates (Figure 6B’, C’) suggest that the relationship between the lifetimes of the network and of individual components tunes force generation by impacting asymmetric component distributions along F-actin. Thus, our parametrization of Δ*α* (Figure 4) likely quantifies structural differences in the filament network architecture among ring types, including component protein abundance and dynamics, and F-actin filament polymerization dynamics.

### Physical modeling and observations of ring closure motivate testable hypotheses of ring contractility

Our calculated inferred turnover rates for NMMII and anillin (Figure 6) indicated that the retention of these structural components is differentially regulated among the subcellular contexts. This motivates several measurements including that of F-actin abundance throughout ring closure, and the turnover (such as fluorescence recovery after photobleaching (FRAP)) of conserved cytoskeletal proteins. Interestingly, our work indicated that the GSC cytokinetic ring generated more active stress than either of the rachis bridges. Laser ablation of rings can be used to measure recoil velocity and compare the relative magnitude of ring intrinsic forces during closure among ring types. In addition, our modeling work indicated that NMMII motoring contributed more to the closure of the GSC cytokinetic ring than to the closure of rachis bridges. Genetically tuning NMMII motor activity (*e*.*g*. via the use of a mutant *C. elegans* strain expressing temperature-sensitive NMMII (Pang *et al*. 2004; Nakamura *et al*. 2005)) could help test whether GSC cytokinesis is more sensitive to temperature shifting than apoptotic and cellularization ring closure.

In sum, our theoretical model implementing *in vivo* measurements reveals that different rings in a syncytium close via the same mechanics but with distinct regulation of protein component abundance, dynamics, and distribution. Our framework makes many testable hypotheses about how protein behavior and abundance dynamics, material properties, and the kinetics of actomyosin ring closure are linked.

## Acknowledgements

We thank Mary Elting, Reem Hakeem, and all members of the AS Maddox, PS Maddox, Fürthauer, and Elting labs for fruitful discussions and helpful reading of the manuscript. We thank Emily Bartle for help with image registration method. SF and AZ have been funded by the Vienna Science and Technology Fund (WWTF)[10.47379/VRG20002]. JBL was supported in part by a grant from NIGMS under award T32 GM119999. This study was also supported by the NIGMS of the National Institutes of Health (R35GM144238), and by the National Science Foundation (2153790) to ASM.

## Author Contributions

ASM, PSM, MEW, BH, JBL conceived and designed experiments and methods. JBL performed experiments and analyzed data, BH and MEW provided technical support for those experiments and analysis. AZ, JBL, and SF formulated model. JBL, AZ, MEW, BH, PSM, SF, and ASM contributed to writing and editing of the manuscript.

