## appendix for "Three types of actomyosin rings within a common cytoplasm exhibit distinct modes of contractility"

### Theory of highly crosslinked cytoskeletal networks

For the coarse-grained description of actomyosin rings in terms of their microscopic constituents we use an active gel theory derived for highly crosslinked networks in (Fürthauer *et al.*, 2021) (Fürthauer *et al.*, 2019). Within this theory the actin filaments are described as rods of length $L$. A filament $i$ is characterized by its center of mass position $\vec{x}_i$ and its orientation $\vec{p}_i$, whereas the position on the filament is parametrized with the parameter $s_i \in [-L/2, L/2]$ (Figure 3A).

For a number of passive crosslinkers $c^x(s_i, s_j)$, bound between points $s_i$ on filament $i$ and point $s_j$ on filament $j$ , the effective force $f_{ij}^x$ exerted between the filaments is assumed to be dominated by friction, i.e. It is taken to be proportional to the velocity difference between attachment points $s_i$ and $s_j$

$$\vec{f}_{ij}^{x} = -\gamma^x c^x(s_i, s_j)(\vec{v}_i + s_i\dot{\vec{p}}_i - \vec{v}_j - s_j\dot{\vec{p}}_j), \qquad \text{(A1)}$$

where $\gamma^x$ is the friction constant for crosslinkers, and $\vec{v}_i = \dot{\vec{x}}_i, \vec{v}_j = \dot{\vec{x}}_{\mathrm{j}}$ describe the center of mass velocity of the filaments. In the case of a motor abundance $c^m(s_i, s_j)$, apart from the frictional coupling of filaments, we expect also the presence of active forces due to the motor stepping. Thus, the exerted force $f_{ij}^m$ between filament $i$ and $j$ in the simplest case of a linear force-velocity curve for the motor, will read

$$\vec{f}_{ij}^{m} = -\gamma^m c^m(s_i, s_j)(\vec{v}_i + s_i\dot{\vec{p}}_i - \vec{v}_j - s_j\dot{\vec{p}}_j + V_{||}(\vec{p}_i - \vec{p}_j)), \qquad \text{(A2)}$$

with $\gamma^m$ standing for the friction constants for motors and $V_{||}$ being the filament sliding speed due to the motor.

To allow for a non-uniform distribution of crosslinkers and motors we postulate that their densities change linearly along the filament length $L$ as follows

$$c^x(s_i, s_j) = c_0^x + c_1^x(s_i + s_j) \qquad \text{(A3)}$$

$$c^m(s_i, s_j) = c_0^m + c_1^m(s_i + s_j). \quad \text{(A4)}$$

In the above expressions, $c_0^{x,m}$ is the total amount of crosslinker or motor per filament, while $c_1^{x,m}$ stand for the gradients of crosslinker and motor distributions along the length of the filament (Figure 3B). For simplicity we assume in this work that the concentration gradient of crosslinkers or motors is directly proportional to their total abundance $c_0^{x,m}$at each time, i.e.

$$c_1^{x,m} = \alpha^{x,m}\, c_0^{x,m}/L, \text{ (A5)}$$

with the proportionality parameters $\alpha^{x,m}$ quantifying the spatial asymmetry of the crosslinking and motor proteins 'distribution along the filament length (Figure 5B).

The total force $\vec{F}_{ij}$ acting from filament $j$ on filament $i$ is then given by the expression

$$\vec{F}_{ij} = \int_{-L/2}^{L/2} ds_i \int_{-L/2}^{L/2} ds_j \int_{\Omega(\vec{x}_i)} d^3\vec{y}\ \delta(\vec{y} - \vec{x}_j - s_j\,\vec{p}_j + s_i\,\vec{p}_i)\big[\big(\vec{f}_{ij}^{x} + \vec{f}_{ij}^{m}\big)\cdot \vec{w}_{ij}\big]\widehat{w}_{ij}, \text{ (A6)}$$

where we have integrated over both filament lengths and a spherical domain $\Omega(\vec{x}_i)$ around $\vec{x}_i$ of radius $R$, indicative of the range of crosslinker and motors interactions. The crosslinker interaction forces $\vec{f}_{ij}^{x,m}$ are projected along the crosslinker direction

$$\widehat{w}_{ij} = \frac{\vec{x}_i+s_i\,\vec{p}_i-\vec{x}_j-s_j\,\vec{p}_j}{\|\vec{x}_i+s_i\,\vec{p}_i-\vec{x}_j-s_j\,\vec{p}_j\|} \quad \text{(A7)}$$

so that the crosslinkers are torque free.

We next introduce the following fields

$$\rho(\vec{x}) = \sum_i \delta\,(\vec{x}_i - \vec{x}) \quad \text{(A8)}$$
$$\vec{v}(\vec{x}) = \frac{1}{\rho(\vec{x})}\sum_i \vec{v}_i\delta\,(\vec{x}_i - \vec{x}) \quad \text{(A9)}$$
$$\vec{P}(\vec{x}) = \frac{1}{\rho(\vec{x})}\sum_i \vec{p}_i\delta\,(\vec{x}_i - \vec{x}) \quad \text{(A10)}$$
$$\boldsymbol{Q}(\vec{x}) = \frac{1}{\rho(\vec{x})}\sum_i \vec{p}_i\vec{p}_i\delta\,(\vec{x}_i - \vec{x}) \quad \text{(A11)}$$
$$\boldsymbol{\mathcal{T}}(\vec{x}) = \frac{1}{\rho(\vec{x})}\sum_i \vec{p}_i\vec{p}_i\vec{p}_i\delta\,(\vec{x}_i - \vec{x}) \quad \text{(A12)}$$
$$\boldsymbol{\mathcal{S}}(\vec{x}) = \frac{1}{\rho(\vec{x})}\sum_i \vec{p}_i\vec{p}_i\vec{p}_i\vec{p}_i\delta\,(\vec{x}_i - \vec{x}), \quad \text{(A13)}$$

representing the filament density $\rho(\vec{x})$, the filament velocity field $\vec{v}(\vec{x})$, the filament polarization $\vec{P}(\vec{x})$, the nematic order $Q(\vec{x})$, the third rank $\mathcal{T}(\vec{x})$ and the fourth rank orientation $\mathcal{S}(\vec{x})$ tensors. The network stress $\boldsymbol{\Sigma}(\vec{x})$ can be derived in terms of the fields (A8)-(A13) using the relation

$$\nabla\cdot\boldsymbol{\Sigma}(\vec{x}) = \sum_{ij} \delta\,(\vec{x}_i - \vec{x})\,\vec{F}_{ij} \quad \text{(A14)}$$

together equation (A6) for the force $\vec{F}_{ij}$ and following the coarse graining procedure in (Fürthauer *et al.*, 2021). The resulting expression for the stress tensor in the apolar case $\vec{P}(\vec{x}) = 0, \mathcal{T}(\vec{x}) = 0,$ is then

$$\boldsymbol{\Sigma}(\vec{x}) = \varrho^2(\vec{x})\eta^{eff}:\nabla\vec{v}(\vec{x}) + \varrho^2(\vec{x})\zeta_1\boldsymbol{Q}(\vec{x}). \quad \text{(A15)}$$

In the above expression, the first term is a viscous term $\boldsymbol{\Sigma}^V$, with the effective viscosity rank-4 tensor $\eta^{eff}_{\alpha\beta\gamma\delta} = \frac{4\pi(\gamma^x c_0^x+\gamma^m c_0^m)}{3}\left\{\frac{L^2R^5}{50}\left[\delta_{\alpha\gamma}\delta_{\beta\delta}+\delta_{\alpha\delta}\delta_{\beta\gamma}+\delta_{\alpha\beta}\delta_{\gamma\delta}\right]+\frac{L^4R^3}{36}\mathcal{S}_{\alpha\beta\gamma\delta}\right\}$ with $\delta_{\alpha\beta}$ the Kronecker delta and the notation $(\boldsymbol{A}:\boldsymbol{B})_{\alpha\beta} = A_{\alpha\beta\gamma\delta}B_{\gamma\delta}$ standing for contraction of two indices, with the repeated index being contracted (Einstein summation convention). The second term is the active term $\boldsymbol{\Sigma}^A$ , proportional to the nematic tensor $\boldsymbol{Q}(\vec{x})$ with $\zeta_1 = -\frac{4\pi L^4R^3}{108}\gamma^m\gamma^x V_{||}\frac{c_1^m c_0^x - c_1^x c_0^m}{(\gamma^x c_0^x+\gamma^m c_0^m)^2}$

To proceed further we need to have a knowledge of the orientation of the actin filaments. Two apolar cases are of particular interest for the description of actomyosin rings: the isotropic and the nematically aligned along the direction $\hat{e}_s$ of the ring's circumference.

**Isotropic case**

In the isotropic 3D case the nematic tensor $Q(\vec{x})$ and the rank-4 $\mathcal{S}(\vec{x})$ tensor is given by

$$Q_{\alpha\beta}(\vec{x}) = \frac{1}{3}\delta_{\alpha\beta} \text{ and } \mathcal{S}_{\alpha\beta\gamma\delta}(\vec{x}) = \frac{1}{15}\left[\delta_{\alpha\gamma}\delta_{\beta\delta}+\delta_{\alpha\delta}\delta_{\beta\gamma}+\delta_{\alpha\beta}\delta_{\gamma\delta}\right]. \quad \text{(A16)}$$

Substituting these in Eq. (A15) and using also (A5) we obtain

$$\boldsymbol{\Sigma}(\vec{x}) = \boldsymbol{\Sigma}^V(\vec{x}) + \boldsymbol{\Sigma}^A(\vec{x}), \quad \text{(A17)}$$

where the viscous term is

$$\boldsymbol{\Sigma}^V(\vec{x}) = \eta_{\square}^{iso}[\nabla\vec{v} + (\nabla\vec{v})^T + (\nabla\cdot\vec{v})\boldsymbol{I}], \quad \text{(A18)}$$

with

$$\eta_{\square}^{iso} = \frac{4\pi R^3 L^2}{30}\left(\frac{L^2}{5} + \frac{R^2}{54}\right)\varrho^2(\vec{x})(\gamma^x c_0^x + \gamma^m c_0^m) \quad \text{(A19)}$$

and $I$ the identity matrix, while the active term reads

$$\boldsymbol{\Sigma}^A(\vec{x}) = -\zeta_{\square}^{iso}\varrho^2(\vec{x})\boldsymbol{I} \quad \text{(A20)}$$

with

$$\zeta_{\square}^{iso} = \frac{1}{3}\frac{4\pi L^3 R^3}{108}\gamma^m\gamma^x V_{||}\frac{(\alpha^m-\alpha^x)c_0^x c_0^m}{\gamma^m c_0^m + \gamma^x c_0^x}. \quad \text{(A21)}$$

**Nematic case**

In the case where the actin filaments are aligned along the direction $\hat{e}_s$ of the ring's circumference (see Fig. 5A) both the nematic tensor $Q(\vec{x})$ and the rank-4 tensor $\mathcal{S}(\vec{x})$ have all entries zero, except for the following

$$Q_{ss}(\vec{x}) = 1 \text{ and } \mathcal{S}_{ssss}(\vec{x}) = 1. \quad \text{(A22)}$$

Then the viscous stress reads

$$\boldsymbol{\Sigma}^V(\vec{x}) = \eta_1^{nem}[\nabla\vec{v} + (\nabla\vec{v})^T + (\nabla\cdot\vec{v})\boldsymbol{I}] + \eta_2^{nem}\nabla_s v_s \boldsymbol{Q}, \quad \text{(A23)}$$

with

$$\eta_1^{nem} = \frac{4\pi R^5 L^2}{150}\varrho^2(\vec{x})(\gamma^x c_0^x + \gamma^m c_0^m) \quad \text{and} \quad \eta_2^{nem} = \frac{4\pi R^3 L^4}{108}\varrho^2(\vec{x})(\gamma^x c_0^x + \gamma^m c_0^m), \quad \text{(A24)}$$

while the active stress is

$$\boldsymbol{\Sigma}^A(\vec{x}) = -\zeta_{\square}^{nem}\varrho^2(\vec{x})\boldsymbol{Q} \quad \text{(A25)}$$

With

$$\zeta_{\square}^{nem} = \frac{4\pi L^3 R^3}{108}\gamma^m\gamma^x V_{||}\frac{(\alpha^m-\alpha^x)c_0^x c_0^m}{\gamma^m c_0^m + \gamma^x c_0^x}. \quad \text{(A26)}$$

**Thin-ribbon approximation**

In order to apply the above active gel theory to actomyosin rings, we need to explicitly account for the ring geometry. For this we introduce a parametrization of the ring by the angle $\theta$, such that the arc length $s$ is $s = r\theta$; where r is the ring radius. We assume that the cytoskeletal network is a thin ribbon with thickness $h$ (Figure 5A). Next, we define the normal vector $\hat{e}_z$ and the tangent vector $\hat{e}_s$ (Figure 5A). Following the approach in (Foster *et al.*, 2022) we can write the force balance $\nabla\cdot\boldsymbol{\Sigma} = 0$ for this geometry given the thin shell approximation $h \ll r$ as

$$\eta_s\frac{1}{r^2}\left(\partial_\theta^2 V^s + \partial_\theta V^z\right) + \frac{1}{r}\partial_\theta\Sigma^A - \gamma V^s = 0 \qquad \text{(A27)}$$

for the tangential direction $\hat{e}_s$ and

$$\frac{\eta_s}{r^2}(\partial_\theta V^s + V^z) + \frac{1}{r}\Sigma^A = 0 \qquad \text{(A28)}$$

for the radial direction $\hat{e}_z$ . In the above expression $V^s$ is the tangential velocity along the ring circumference opposed by external friction $\gamma$ due to interactions with the plasma membrane and actomyosin cortex, while $V^z$ is the normal velocity which induces changes in the radius $r$. The viscosity $\eta_s$ is the viscosity coefficient in the term $\Sigma_{ss}^V$ , while $\Sigma^A$ stands for $\Sigma_{ss}^A$ . Given the expressions (A17)-(A26) these can be written as

$$\eta_s^{\square} = b_1\varrho^2(\vec{x})(\gamma^x c_0^x + \gamma^m c_0^m) \quad \text{(A29)}$$

with

$$b_1 = \frac{4\pi R^3 L^2}{15}\left(\frac{L^2}{5} + \nu\frac{R^2}{54}\right)\varrho^2(\vec{x})(\gamma^x c_0^x + \gamma^m c_0^m) \quad \text{(A30)}$$

and

$$\Sigma^A = -\, b_2\varrho^2(\vec{x})\gamma^m\gamma^x V_{||}\frac{(\alpha^m - \alpha^x)c_0^x c_0^m}{\gamma^m c_0^m + \gamma^x c_0^x} \quad \text{(A31)}$$

with

$$b_2 = \frac{4\pi L^3 R^3}{108}\lambda. \quad \text{(A32)}$$

In the above equations the parameters $\nu$ and $\lambda$ have different values for the different orientations of actin filaments. According to the above discussion $\nu = 1$ and $\lambda = \frac{1}{3}$ for the isotropic case, while $\nu = 15$ and $\lambda = 1$ for the nematic one. By Fourier transforming Eqs. (A27), (A28) and solving for the normal velocity $V^z$ we can calculate (Foster et al., 2022) the contraction rate of the actomyosin radius $r$ as

$$\frac{dr}{dt} = -r\frac{\Sigma^A}{\eta_s}, \qquad \text{(A33)}$$

or, given Eqs. (A29)-(A31) as

$$\frac{dr}{dt} = A\gamma^m\gamma^x V_{||} r\frac{(\alpha^m - \alpha^x)c_0^x c_0^m}{(\gamma^m c_0^m + \gamma^x c_0^x)^2} \quad \text{(A34)}$$

with

$$A = \frac{5L\lambda}{36\left(\frac{L^2}{5} + \nu\frac{R^2}{54}\right)}, \quad \text{(A35)}$$

and $\frac{1}{3} \le \lambda \le 1$, $1 \le \nu \le 15$, as discussed above.

Given the expression

$$c_0^{x,m}(t) = \beta^{x,m}\frac{I^{x,m}(t)}{2\pi r} \qquad \text{(A36)}$$

Relating the concentrations $c_0^{x,m}(t)$ to corresponding fluorescence intensities $I^{x,m}(t)$ and using $\Delta\alpha = AV_{||}(\alpha^x - \alpha^m)$, as well as $\beta = \frac{\beta^m\gamma^m}{\beta^x\gamma^x}$ we arrive at

$$\frac{dr}{dt} = -\beta \cdot \Delta\alpha \cdot r\frac{I^m(t)I^x(t)}{(\beta I^m(t) + I^x(t))^2}. \quad \text{(A37)}$$

**Mean contraction rate**

From the above we have derived the contraction rate of actomyosin rings in terms of motors and crosslinker densities or fluorescence intensities (Eqs. (A34),(A37)) and other microscopic details. We would like now to link this to the average closure speed examined in (Carvalho *et al.*, 2009; Calvert *et al.*, 2011; Bourdages *et al.*, 2014).

For this we must solve Eq. (A37), resulting in

$$r(t) = r(0)\exp\left[-\beta\Delta\alpha\int_0^t \frac{I^m(t')I^x(t')}{(\beta I^m(t') + I^x(t'))^2}dt'\right]. \quad \text{(A38)}$$

Then we can extract the average rate

$$\frac{\Delta C}{\Delta t} = \frac{C(0) - C(T)}{T} = 2\pi\frac{r(0) - r(T)}{T} = C(0)\frac{1}{T}\left\{1 - \exp\left[-\beta\Delta\alpha\int_0^T \frac{I^m(t')I^x(t')}{(\beta I^m(t') + I^x(t'))^2}dt'\right]\right\}. \quad \text{(A39)}$$

Comparing with the scaling law $\frac{\Delta C}{\Delta t} = mC(0) + b$ we can identify

$$m = \frac{1}{T}\left\{1 - \exp\left[-\beta\Delta\alpha \int_0^T \frac{I^m(t\prime)I^x(t\prime)}{\left(\beta I^m(t\prime)+I^x(t\prime)\right)^2} dt'\right]\right\}, \text{(A40)}$$

or for short times $T$

$$m = \beta\Delta\alpha \left\langle\frac{I^m(t\prime)I^x(t\prime)}{\left(\beta I^m(t\prime)+I^x(t\prime)\right)^2}\right\rangle. \text{(A41)}$$

with the average $\langle \square \rangle$ implying an average in time from 0 to $T$. For an identification of coefficient $b$ another force opposing constrictions should be added to the above model, but this goes beyond the scope of the current study.
